# Novel integrated computational AMP discovery approaches highlight diversity in the helminth AMP repertoire

**DOI:** 10.1101/2023.02.02.526830

**Authors:** Allister Irvine, Darrin Mckenzie, Ciaran J. McCoy, Robert Graham, Ciaren Graham, Sharon A. Huws, Louise E. Atkinson, Angela Mousley

## Abstract

Antimicrobial Peptides (AMPs) are immune effectors that are key components of the invertebrate innate immune system providing protection against pathogenic microbes. Parasitic helminths share complex interactions with their hosts and closely associated microbiota that are likely regulated by a diverse portfolio of antimicrobial immune effectors including AMPs. Knowledge of helminth AMPs has largely been derived from nematodes, whereas the flatworm AMP repertoire has not been described.

This study highlights limitations in the homology-based approaches, used to identify putative nematode AMPs, for the characterisation of flatworm AMPs, and reveals that innovative algorithmic AMP prediction approaches provide an alternative strategy for novel helminth AMP discovery. The data presented here: (i) reveal that flatworms do not encode traditional lophotrochozoan AMP groups (Big Defensin, CSαβ peptides and Myticalin); (ii) describe a unique integrated computational pipeline for the discovery of novel helminth AMPs; (iii) reveal >16,000 putative AMP-like peptides across 127 helminth species; (iv) highlight that cysteine-rich peptides dominate helminth AMP-like peptide profiles; (v) uncover eight novel helminth AMP-like peptides with diverse antibacterial activities, and (vi) demonstrate the detection of AMP-like peptides from helminth biofluids. These data represent a significant advance in our understanding of the putative helminth AMP repertoire and underscore a potential untapped source of antimicrobial diversity which may provide opportunities for the discovery of novel antimicrobials. Further, unravelling the role of endogenous worm-derived antimicrobials and their potential to influence host-worm-microbiome interactions may be exploited for the development of unique helminth control approaches.

**Author summary:** Invertebrate antimicrobial peptides (AMPs) form the first line of defence against pathogenic microbes. Helminths are worms (flatworm, roundworm) that live in microbe-rich environments throughout their lifecycles however little is known about how they protect themselves against pathogens or how they interact with microbes. Understanding AMP profiles in helminths, their importance to helminth biology, and how they shape microbial communities could reveal novel approaches for anthelmintic and/or antimicrobial development.

In this study we describe a novel integrated homology- and computational-based pipeline for the discovery of helminth AMPs. This approach revealed that, whilst flatworms do not possess traditional AMPs, they have a repertoire of unique AMP-like peptides that are predominantly cysteine-rich. Significantly eight novel helminth AMP-like peptides, discovered using this pipeline, have antibacterial activities against a range of bacteria highlighting their potential as novel antimicrobials. Further, peptidomics analyses demonstrate the presence of AMP-like peptides in helminth body fluids supporting the need to further characterise these peptides and their function(s) in helminths. These data present novel opportunities to better understand helminth biology, discover new antimicrobials and develop future control strategies for helminth parasites.

## Introduction

Antimicrobial peptides (AMPs) are ubiquitous natural immune components that play key roles in protection against microbial threat [1]. In response to mounting antimicrobial resistance (AMR) pressures, AMPs have been highlighted as potential alternatives to antibiotics due to their broad-spectrum antimicrobial activities and structural diversity [2]. Early AMP research relied on the biochemical isolation of bioactive peptides. These approaches were costly and limited to organisms which are easily accessible and readily maintained *in vitro* [3], such that >50% of the AMP cohort on the Antimicrobial Peptide Database (APD3) is derived from arthropods and amphibians [4]. Advances in sequencing technologies coupled with a rapid rise in the availability of omics data have since enabled the identification of AMPs based on sequence homology across diverse species [3]. Whilst successful, these approaches limit the identification of AMPs to those that have well characterised motifs and hinder the discovery of novel AMP families. Over the last decade, novel approaches to AMP discovery have been adopted including the use of machine learning algorithms [5]. Seeding machine learning algorithms with known AMP datasets has driven the development of AMP prediction tools which can identify putative AMPs from complex omics datasets. Many of these prediction tools are trained to recognise the physicochemical properties of AMPs, in addition to AMP sequence motifs, allowing the identification of novel peptides [6]. Indeed, these computational approaches have successfully identified novel AMPs from diverse organisms including the cuttlefish (*Sepia officinalis*) [7], the fly (*Hermetia illucens*) [8], and the human gut microbiome [9].

Parasitic helminths live in diverse host niches where they are exposed to a wide range of microbes, such that the production of a diverse portfolio of helminth-derived antimicrobials could enable them to shape their immediate microbial environment. Interestingly, parasitic helminths, including cestodes, trematodes and nematodes, are known to modify host microbiota [10-15]. In other invertebrates, AMPs are important in establishing and maintaining tissue-specific microbiomes including the *Drosophila melanogaster* gut microbiome [16] and the *Hydra vulgaris* embryonic microbiome [17].

Recent studies highlight that phylum Nematoda is AMP-rich [18, 19], however equivalent data for members of the phylum Platyhelminthes are not available beyond *Schistosoma mansoni* [20, 21]. Moreover, the homology-based approach employed in the nematode studies did not facilitate the discovery of putative AMPs that are non-traditional and sequence divergent. This study aims to explore the flatworm AMP repertoire and address the limitations of homology-directed approaches for AMP discovery through the development of an innovative computational pipeline to aid discovery of novel AMPs within nematodes and flatworms. The data presented here: (i) reveal that flatworms do not encode traditional lophotrochozoan AMP groups (Big Defensin, CSαβ peptides and Myticalin); (ii) describe a unique integrated computational pipeline for the discovery of novel helminth AMPs; (iii) reveal >16,000 AMP-like peptides across 127 helminth species; (iv) highlight that cysteine-rich peptides dominate helminth AMP-like peptide profiles; (v) uncover eight novel helminth AMP-like peptides with diverse antibacterial activities, and (vi) demonstrate the detection AMP-like peptides in helminth biofluids.

To our knowledge, these data represent the first phylum-spanning computational approach to novel AMP discovery in helminths and provide a route to unravelling the unique characteristics of flatworm and nematode AMPs that do not share sequence similarity to other known invertebrate AMPs. The data generated in this study will be valuable for future helminth AMP discovery efforts and will provide a springboard for functional biology and novel therapeutic discovery.

## Materials and Methods

### Construction of a lophotrochozoan AMP library

To construct a comprehensive lophotrochozoan-derived AMP library, published literature was searched for known AMPs from 15 lophotrochozoan phyla (Annelida, Brachiopoda, Bryozoa, Chaetognatha, Cycliophora, Dicyemida, Entoprocta, Gastrotricha, Gnathostomulida, Micrognathozoa, Mollusca, Nemertea, Phoronida, Platyhelminthes and Rotifera) as indicated by Bleidorn (22). In addition, AMP databases (see Table 1 and Fig 1A) were also mined for naturally derived lophotrochozoan AMPs but any modified or artificially designed AMPs from lophotrochozoan species were not included in the library. Lophotrochozoan-derived AMPs were categorised into AMP groups based on homology and/or the presence of AMP motifs.

**Table 1:** AMP databases used to identify lophotrochozoan-derived AMPs.

| Database name | URL | Reference |
| --- | --- | --- |
| Antimicrobial Peptide Database (APD3) | https://aps.unmc.edu/home | [23] |
| Collection of Antimicrobial Peptides (CAMP<sub>R3</sub>) | www.camp.bicnirrh.res.in/ | [24] |
| Yet Another Database of Antimicrobial Peptides (YADAMP) | www.yadamp.unisa.it/ | [25] |
| Database of Antimicrobial Activity and Structure of Peptides (DBAASP<sub>v3.0</sub>) | www.dbaasp.org/ | [26] |
| Invertebrate Antimicrobial Peptide Database (InverPep) | https://ciencias.medellin.unal.edu.co/gruposdeinvestigacion/prospeccionydisenobiomoleculas/InverPep/public/home_en | [27] |
| Data Repository of Antimicrobial Peptides (DRAMP) | http://dramp.cpu-bioinform.org/ | [28] |

**Fig 1:**
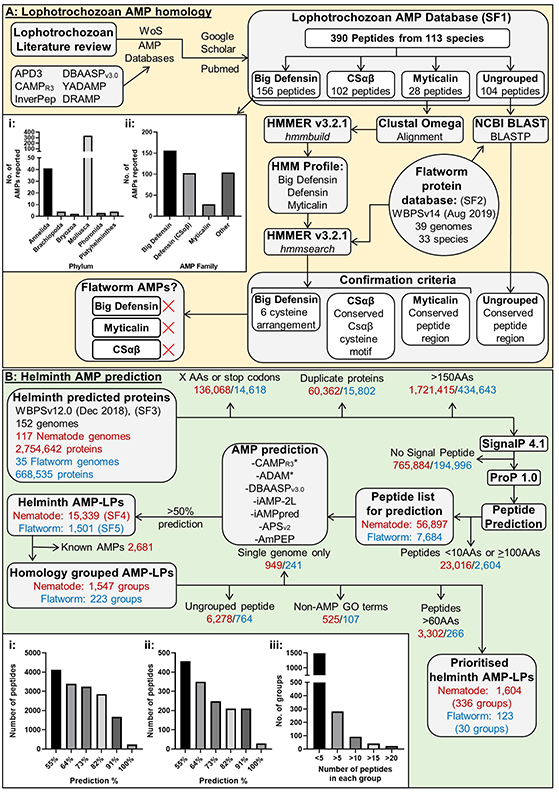
Integrated computational AMP discovery pipeline reveals novel helminth AMP-like peptides. A) Summary of homology directed approach using lophotrochozoan-derived AMPs. i) Breakdown of lophotrochozoan AMP database by phylum. ii) Breakdown of lophotrochozoan AMP database by AMP family. Asterisk indicates that multiple algorithms were used for each AMP prediction tool. B) Summary of computational helminth AMP prediction approach. i) The number of nematode AMP-LPs and percentage of prediction tools indicating antimicrobial potential. ii) The number of flatworm AMP-LPs and percentage of prediction tools indicating antimicrobial potential. iii) The number of peptides in each group after AMP-LPs were organised into groups. AA, Amino acid; ADAM, A Database of Antimicrobial Peptides; AMP, Antimicrobial Peptide; AMP-LP, Antimicrobial Peptide-like Peptide; APD, Antimicrobial Peptide Database; APS, Antimicrobial Peptide Scanner; BLAST, Basic Local Alignment Search Tool; BLASTP, Protein BLAST; CAMP, Collection of Antimicrobial Peptides; CSαβ, Cysteine stabilised α-helix and β-sheet fold peptides; Cys, Cysteine; DBAASP, Database of Antimicrobial Activity and Structure of Peptides; DRAMP, Data Repository of Antimicrobial Peptides; GO, Gene Ontogeny; GREP, Globally search a Regular Expression and Print; HMM, Hidden Markov Model; iAMPpred, Improved Prediction of Antimicrobial Peptides; InverPep, Invertebrate Antimicrobial Peptide Database; NCBI, National Center for Biotechnology Information; nr, Non-redundant protein sequence database; SF, Supplementary File; TBLASTN, Translated Nucleotide BLAST; WBPS, WormBase Parasite; WoS, Web of Science; YADAMP, Yet Another Database of Antimicrobial Peptides.

### Identification of flatworm AMP-encoding genes using homology-directed approaches

Putative flatworm AMP-encoding genes were identified using Basic Local Alignment Search Tool (BLAST) and Hidden Markov Model (HMM) homology searches as previously described [19]. Briefly, for HMM analyses a multiple sequence alignment (MSA) of each lophotrochozoan AMP group (Big Defensin, Cysteine stabilised α-helix and β-sheet fold peptides [CSαβ] and Myticalin, see Supplementary File 1) was generated using Clustal Omega [29]. Where appropriate, alignments were modified to ensure correct alignment of AMP sequence motifs and group-specific HMM profiles were built using the *hmmbuild* command in HMMER v3.2.1 (www.hmmer.org). A database of flatworm predicted proteins (33 flatworm species; see Supplementary File 2) corresponding to Wormbase Parasite version 14 [WBPS; 30] was concatenated and the *hmmsearch* command was used to search the HMM profiles against this database. All HMM returns (E value <10) were manually assessed for AMP motifs.

Lophotrochozoan AMPs that do not belong to known AMP groups were not suitable for HMM analyses and therefore were employed as queries in BLASTP searches against the flatworm predicted protein database (33 flatworm species; see Supplementary File 2) using NCBI BLAST version 2.10.0+ (standard settings). BLASTP returns (E value <10) were assessed for the presence of specific AMP motifs or high peptide conservation. If putative peptides were not identified via BLASTP, Translated Nucleotide BLAST (TBLASTN) searches were conducted using NCBI BLAST version 2.10.0+ (standard settings) against a concatenated genome dataset for the 33 flatworm species (obtained from WBPSv14). This approach ensures that unannotated genes are not overlooked.

### Identification of novel helminth AMP-encoding genes using the computational AMP prediction pipeline

#### Datasets analysed and peptide identification approach

Predicted protein datasets from 96 nematode and 31 flatworm species (152 genomes in total; 117 nematode and 35 flatworm genomes; see Fig 1B and Supplementary File 3) were obtained from WBPS version 12.0 [30]. Duplicate protein sequences and proteins containing unknown amino acids (denoted X) or stop codons (denoted *) were removed from each predicted protein dataset. Proteins >150AAs in length were removed and those remaining were analysed for the presence of a signal peptide using SignalP 4.1 [https://services.healthtech.dtu.dk/service.php?SignalP-4.1; Eukaryotes Organism Group, Default D-cutoff values; 31]. Proteins with signal peptides were analysed using ProP 1.0 [https://services.healthtech.dtu.dk/service.php?ProP-1.0; Standard Settings; 32] to predict propeptide cleavage sites. Putative peptides were predicted based on the positions of the signal peptide and propeptide cleavage sites. All potential peptides were included where multiple cleavage sites were predicted in a single protein. Predicted peptides outside of the recommended lengths for AMP prediction (<10 and ≥100 amino acids) were removed from the analysis pipeline.

#### AMP prediction using computational tools

The helminth predicted peptides (see Fig 1B) were analysed using 11 computational AMP prediction tools (see Table 2). These tools were chosen based on: (i) reported sensitivity and specificity for AMP prediction, (ii) evidence of successful use in other studies, (iii) amenability to high-throughput sequence input and (iv) the ability to predict general antimicrobial activity rather than activity against a specific microbe [24, 26, 33-37]. Each prediction tool was applied using standard settings via web-based platforms with the exception of AmPEP which was hosted locally on MATLAB version 9.5 (R2018b). A peptide sequence was designated as an AMP-like peptide (AMP-LP) if antimicrobial activity was indicated in >50% of the prediction tools employed.

**Table 2:** AMP prediction tools employed in this study.

| **Algorithm:** | **Website:** | **Reference:** |
| --- | --- | --- |
| CAMP<sub>R3</sub> SVM | http://www.camp.bicnirrh.res.in/predict/ | [24] |
| CAMP<sub>R3</sub> RF | http://www.camp.bicnirrh.res.in/predict/ | [24] |
| CAMP<sub>R3</sub> ANN | http://www.camp.bicnirrh.res.in/predict/ | [24] |
| CAMP<sub>R3</sub> DA | http://www.camp.bicnirrh.res.in/predict/ | [24] |
| ADAM SVM | http://bioinformatics.cs.ntou.edu.tw/ADAM/svm_tool.html | [33] |
| ADAM HMM | http://bioinformatics.cs.ntou.edu.tw/ADAM/hmm_tool.html | [33] |
| DBAASP<sub>v3.0</sub> | https://dbaasp.org/tools?page=general-prediction | [26] |
| iAMP-2L | http://www.jci-bioinfo.cn/bioinfo/iAMP-2L | [34] |
| iAMPpred | http://cabgrid.res.in:8080/amppred/ | [35] |
| APS<sub>vr.2</sub> | https://www.dveltri.com/ascan/v2/ascan.html | [36] |
| AmPEP | https://sourceforge.net/projects/axpep/ | [37] |

ADAM, A Database of Anti-Microbial Peptides; APS_vr.2_, Antimicrobial Peptide Scanner (Feb2019 model); ANN, Artificial Neural Network; CAMP_R3_, Collection of Anti-Microbial Peptides.; DBAASP_v3.0_, Database of Antimicrobial Activity and Structure of Peptides; DA, Discriminant Analysis; HMM, Hidden Markov Model; iAMPpred, Improved Prediction of Antimicrobial Peptides; RF, Random Forest; SVM, Support Vector Machine.

#### Generation of a high-confidence helminth AMP-LP library

Helminth AMP-LPs were filtered and prioritised to generate a library of high-confidence novel peptides. To facilitate the identification of novel AMP-LPs, previously characterised, homology-based nematode AMPs [19] were removed from the pipeline (see Fig 1B). To identify similar peptides within the helminth AMP-LP library, each AMP-LP was used as a BLASTP search query (locally via NCBI-BLAST-2.10.0+) against the helminth AMP-LP library; hits with an overall score >100 were assigned as sequelogs and grouped accordingly (see Supplementary Files 4 and 5). Note that nematode and flatworm AMP-LPs were analysed and grouped separately. To aid data handling and prioritise more broadly conserved peptides, those peptides that did not share homology or originated from a single genome assembly were removed from the pipeline. For the remaining helminth AMP-LPs, Gene Ontogeny [GO; 38, 39] Biological Process and Molecular Function terms were retrieved from WBPSv12; peptide groups which had established non-antimicrobial-related GO terms were removed from the pipeline to reduce likely false positives. Finally, peptide groups, where the majority of peptides were >60 AAs in length, were also removed from the pipeline.

#### Selection of helminth AMP-LPs for antimicrobial screening

Prioritised helminth AMP-LPs, predicted to have linear structure (peptides with <1 cysteine residue), were selected for synthesis and antimicrobial activity screening. Linear helminth AMP-LPs were selected based on the number of AMP prediction tools indicating potential antimicrobial activity and the conservation of the peptides across multiple genes and species. Where there were peptide variations within an AMP-LP group, the most representative peptide based on peptide conservation to the group consensus sequence was selected for synthesis.

For AMP-LPs selected for synthesis further analyses were conducted in order to characterise the peptides. Full prepropeptide sequences were used as InterPro [https://www.ebi.ac.uk/interpro/; 40] and Pfam [http://pfam.xfam.org/; 41] queries to identify known protein domains or motifs. Where available, relevant RNAseq datasets (analysed and hosted on WBPSv16) were utilised to determine whether AMP-LP-encoding genes were expressed. For this, a Transcripts Per Million (TPM) cut off of two was used to distinguish between expressed and non-expressed genes [42].

### Minimum Inhibitory Concentration assays for helminth AMP-LPs

Prioritised helminth AMP-LPs were synthesised at >70% purity (Genosphere Biotechnologies). Where a peptide possessed a C-terminal glycine residue, it was synthesised with a C-terminal amidation instead of the glycine residue. Peptides and control antibiotics [Ciprofloxacin (for gram-negative species; Sigma-Aldrich) and Vancomycin hydrochloride (for gram-positive species; Sigma-Aldrich)] were dissolved in sterile ultrapure water at a concentration of 5.12mg/ml.

Minimum inhibitory concentrations (MICs) were determined for gram-positive and gram-negative bacterial isolates (see Table 3 for species) using microbroth dilution assays as described previously [43]. Briefly, a single colony from each bacterial species was inoculated in cation-adjusted Mueller Hinton Broth (MHB) and grown for 18-24 hours at 37°C with shaking (225rpm) before being adjusted to a final bacterial density of 5×10^5^ colony forming units (CFU) per ml. Peptides and Vancomycin hydrochloride were diluted using sterile MHB in sterile U-bottom 96 well polypropylene microplates (Greiner Bio One, UK) to give a concentration gradient ranging from 512-0.25μg/ml. Ciprofloxacin was further diluted in sterile MHB to give a concentration gradient ranging from 32-0.02μg/ml. Microplates were incubated at 37°C for 18-24 hours. Plates were assessed for visible growth and the MIC was defined as the lowest concentration of peptide or antibiotic that inhibited visible growth of a bacterial culture after an 18–24-hour incubation. MIC data were validated by positive and negative control antibiotic MIC data for each bacterial isolate [43, 44]. Peptides were considered AMPs if they had an MIC of <100μg/ml which is consistent with the APD3 activity criteria [23].

**Table 3:** Bacterial isolates used to determine Minimum Inhibitory Concentrations of helminth AMP-LPs.

| Bacterial species | Strain | Gram status | Resistance status |
| --- | --- | --- | --- |
| *Escherichia coli* | K12 | Negative |  |
| *Acinetobacter baumannii* | DSMZ 30007 | Negative |  |
| *Pseudomonas aeruginosa* | PA01 | Negative |  |
| *Staphylococcus aureus* | RN4220 | Positive | Methicillin-Sensitive |
| *Staphylococcus aureus* | EMRSA-15 | Positive | Methicillin-resistant |
| *Enterococcus faecalis* | JH2-2 | Positive |  |

### LC-MS/MS detection of AMP-LPs in helminth biofluids

#### Collection and maintenance of *Ascaris suum*

Adult *A. suum* were collected at a local abattoir (Karro Food Group Ltd., Cookstown, Northern Ireland), transported to the laboratory in saline (0.9% NaCl), and maintained in Ascaris Ringers Solution (ARS: 13.14 mM NaCl, 9.47 mM CaCl_2_, 7.83 mM MgCl_2_, 12.09 mM C_4_H_11_NO_3_/ Tris, 99.96 mM NaC_2_H_3_O_2_, 19.64 mM KCl, pH 7.8) at 37 °C until use.

#### Collection of *Ascaris suum* pseudocoelomic fluid

*Ascaris suum* pseudocoelomic fluid (As-PCF) was collected from ~20 female *A. suum* (>20cm) within 3 hours of collection as previously described [45]. A total volume of 10ml As-PCF was collected for each biological replicate (n=3). From each 10ml biological replicate 1ml was transferred to a 2ml low-binding microcentrifuge tube (Eppendorf, UK) and placed on ice prior to LC-MS/MS analysis.

#### Acidified methanol treatment of *As*-PCF

As-PCF was treated with 1ml of ice-cold acidified methanol (Ac-MeOH, 90:9:1 – methanol:ultrapure water (18.2Ω):Acetic acid) as previously described [45] with minor modifications including the use of a glass Dounce homogeniser (Sigma-Aldrich, UK) to resuspend the centrifuged As-PCF pellet. Homogenisation was performed for 60 secs and repeated until the pellet was fully resuspended in solution. Samples were then centrifuged at 19,000g for 15 mins at 4°C and the supernatant was divided across two 2ml low-binding microcentrifuge tubes. 250μl of ultrapure water was added to each tube to reduce the methanol concentration below 60% prior to further processing. Please note that all solvents used for LC/MS preparation and analysis were Optima Grade unless otherwise stated.

#### As-PCF filtering for peptidomic analysis

As-PCF was filtered prior to LC-MS/MS as previously described [45] with minor modifications including an additional 50:50 MeOH:water wash step prior to As-PCF sample loading. In addition, during filtration As-PCF samples were gently pipetted every 20 mins to resuspend any solids that had accumulated in the filter. Finally, the 10kDa flowthrough was then split across two 2ml low-binding microcentrifuge tubes (~1.3ml in each tube) and dried overnight at room temperature using a vacuum concentrator (Eppendorf, UK).

#### As-PCF peptide desalting

As-PCF samples were resuspended in 50μl of 0.1% formic acid via sonication for 3 mins in a benchtop water bath sonicator (Fisher Scientific, UK) and vortexed for 30 secs; this was repeated until the sample was fully resuspended. As-PCF samples were centrifuged at 16,000g for 10 mins to pellet any debris. The supernatant was removed and placed into a fresh 2ml low-binding microcentrifuge tube. Custom STop And Go Extraction (STAGE) tips were produced using established protocols [46]. STAGE tips were pre-treated with 50μl 80% methanol, 0.1% formic acid and centrifuged at 3000rpm for 3 mins (repeated three times). Tips were conditioned through the addition of 50μl 80% acetonitrile, 0.1% formic acid and centrifuged at 3000rpm for 3 mins (repeated three times). STAGE tips were then prepared for sample loading by washing with 50μl 0.1% formic acid and centrifuged at 3000rpm for 3 mins (repeated three times). 50μl of centrifuged As-PCF was loaded into a STAGE tip and centrifuged again at 3000rpm for 3 mins. STAGE tips were then washed with 50μl 0.1% formic acid and centrifuged at 3000rpm for 3 mins (repeated ten times or until the colour was removed from the STAGE tip). After washing, STAGE tips were transferred to a fresh 2ml low-binding microcentrifuge tube and, through the addition of 50μl 80% acetonitrile and 0.1% formic acid and centrifugation at 3000rpm for 3 mins, peptides were eluted.

This was repeated once to ensure all peptides were eluted from the STAGE tip (final volume 100μl). Peptide samples were dried using a vacuum concentrator at room temperature (1-2 hours) and stored at −20°C prior to LC-MS/MS analysis, or at −80°C if LC-MS/MS was delayed for more than 3-5 days.

#### LC-MS/MS analysis

Stored peptide samples were dissolved in 9μl 3% acetonitrile and 0.1% formic acid. Micro-LC-MS/MS analysis was carried out by injection of 8μl of peptide sample into a Eksigent Expert™ Nano LC system (Eksigent, Dublin, Ca) coupled to a Sciex Triple-TOF 6600 mass spectrometer (AB Sciex, Warrington, UK). A Kinetex 2.6μm XB-C18 100 A (150mm x 0.3mm, Phenomenex, UK) column was used for chromatographic separation. Mobile phase A consisted of 100% H_2_O with 0.1% formic acid. Mobile phase B consisted of 100% acetonitrile and 0.1% formic acid. Peptides were separated with a 5μl/min linear gradient of 5-25% B for 68 mins, 35-45% B for 5 mins, 80% B for 3 mins and 3 mins equilibration at 5% B. Data were collected in positive electrospray ionisation (ESI) data-dependant mode (DDA). The 30 most abundant ions were selected for MS/MS following a 250ms TOF-MS survey scan and 50ms MS/MS scan. Dynamic exclusion time was set to 15s. Selected parent ions had charged states between 2 and 4 and were fragmented by Collision-induced dissociation (CID).

#### LC-MS/MS data analysis

MicroLC-ESI-MS raw data were analysed by PEAKS studio X (Bioinformatics solution Inc., Waterloo, ON, Canada). The error tolerances for parent mass and fragment mass were set as 15ppm and 0.1Da, respectively. An enzyme search with unspecific digestion was used. Post-translational modifications were as follows: C-terminal amidation, Pyro glut–Q, Pyro-glut-E, sulfation and oxidation of methionine. A custom prepropeptide AMP library was used in the peptide search process. The peptide database was compiled via comprehensive *in silico* analyses [see above methods and 19]. Peptides were considered as high-confidence positive identifications if detected above the Peptide-Spectrum Match (PSM) 1% FDR cut-off and at least 1 unique peptide was present in at least one biological replicate. Peptides were considered as medium and low confidence if detected above the PSM P-value <0.01 or P-value <0.05 cut offs and at least 1 unique peptide was present in at least one biological replicate. Manual validation was performed on all positively identified peptides to ensure the presence of at least three consecutive b- or y-ions in MS2 spectra were detected.

## Results/Discussion

### Previously identified flatworm AMPs are not conserved across phylum Platyhelminthes

Eight AMPs were previously reported in *Schistosoma mansoni* [20, 21, 47; see Table 4]. In this study, BLAST-based homology analysis did not confirm any of these peptides in *S. mansoni* or reveal conservation across phylum Platyhelminthes, with the exception of *S. mansoni* Schistocins, and a putative Schistocin sequelog in *Schistosoma rodhaini* (WBPS Gene ID: SROB_0000009401). Schistocin peptides have not been biochemically isolated, and the lack of conservation of these peptides and other *S. mansoni* derived AMPs [20, 47] across flatworm species suggests that they do not represent conserved flatworm AMP groups. As a result, it is difficult to direct flatworm AMP discovery through homology-based searches using known flatworm AMPs. This provides a rationale for expanding the homology-based approach to include AMPs derived from other Lophotrochozoa as seed sequences for BLAST searches.

**Table 4:** Reported AMPs from *Schistosoma mansoni*.

| **Peptide Name** | **Reference** | **Activity reported** | **Notes** |
| --- | --- | --- | --- |
| Dermaseptin-like peptide | [20] | Antibacterial<br>Antifungal<br>Haemolytic | Isolated from cercarial extracts |
| Schistocin-1 | [21] | All activity >100µM | Computationally predicted from Kunitz inhibitor protein |
| Schistocin-2 | [21] | Antifungal | Computationally predicted from Kunitz inhibitor protein |
| Schistocin-3 | [21] | Antibacterial<br>Antifungal | Computationally predicted from Kunitz inhibitor protein. Additional peptide designed from this (Schistocin 3.1) |
| N/A | [47] | None | Scorpion Na-channel toxin CSαβ subfamily [GenBank: EX499237] |
| N/A | [47] | None | Scorpion Na-channel toxin CSαβ subfamily [GenBank: EX499261] |
| N/A | [47] | None | Scorpion Na-channel toxin CSαβ subfamily [GenBank: EX499243] |
| N/A | [47] | None | Scorpion Na-channel toxin CSαβ subfamily [GenBank: EX499256] |

### Flatworms do not encode major groups of lophotrochozoan-type AMPs

390 AMPs were identified from 113 lophotrochozoan species (see Supplementary File 1) where molluscan-derived AMPs dominate (see Fig 1Ai). Published literature indicates that lophotrochozoan AMP groups (CSαβ, Big Defensin and Myticalin) are broadly conserved across lophotrochozoan phyla where Big Defensins appear to be the most abundant (see Fig 1Aii). Homology-based searches of available flatworm omics datasets for lophotrochozoan AMPs reveal that members of phylum Platyhelminthes do not encode putative homologs for any of the three major lophotrochozoan AMP groups (CSαβ, Big Defensin and Myticalin). While the absence of Big Defensin-encoding genes in flatworms has been noted previously [48], the absence of CSαβ superfamily peptides is surprising considering CSαβ peptides are widely distributed across diverse taxa (Porifera through Arthropoda) [47].

The absence of canonical lophotrochozoan-type AMPs in flatworms suggests that flatworms may possess AMP groups that are currently undiscovered and distinct from those found in other lophotrochozoan phyla. This finding drives the development and application of alternative approaches for AMP discovery in flatworms.

### A unique integrated computational pipeline reveals novel helminth AMP-like peptides

In addition to the limitations of the homology-based analyses described in this study for flatworms, reduced AMP profiles have been reported in some nematode clades [19]. Indeed, within phylum Nematoda there are major differences in AMP profiles across species despite sharing common microbe-facing environmental and host niches [19]. These observations may suggest the presence of novel AMPs in helminths that are sequence divergent and therefore more difficult to identify using homology-based approaches. This study describes an innovative computational AMP prediction pipeline (see Fig 1B) and its application for the discovery of novel helminth AMPs, providing a springboard for functional biology and novel therapeutic discovery.

#### The peptide prediction phase of the novel AMP prediction pipeline reveals >60,000 putative helminth peptides

The helminth predicted protein datasets employed in this study (2,754,642 nematode and 668,535 flatworm proteins; see Supplementary File 2 and 3) were filtered to remove proteins according to the helminth AMP prediction pipeline (see Fig 1B). Briefly, duplicate protein sequences (60,362 nematode and 15,802 flatworm proteins) and those containing stop codons or uncharacterised amino acids (AAs) (136,068 nematode and 14,618 flatworm proteins; ~7% of all helminth predicted proteins) were removed (see Fig 1B). The remaining proteins were filtered by AA length (proteins >150AAs were removed; 1,721,415 nematode and 434,643 flatworm proteins; ~63% of all helminth predicted proteins). Sequences that were <150AAs were retained based on the length of known AMPs; indeed, ~85% of ecdysozoan and ~98% of amphibian AMP precursors are <150AAs [UniProt Knowledgebase; 49]. Proteins without a putative signal peptide were also removed (765,884 nematode and 194,996 flatworm proteins; ~28% of all helminth predicted proteins). Secretory peptides were predicted from the remaining cohort of predicted proteins (70,913 nematode and 8,476 flatworm proteins; 2% of the original helminth predicted proteins) based on the position of the putative signal peptide and any putative propeptide cleavage sites. At this stage 79,389 proteins remained which were predicted to encode 90,201 peptides. At this stage any peptides <10AAs or >100AAs in length were also removed (23,016 nematode and 2,604 flatworm peptides) as these fall outside the parameters of the AMP prediction tools employed here (see Table 2). A final cohort of 64,581 helminth predicted peptides (56,897 nematode and 7,684 flatworm peptides) were channelled into the computational AMP prediction tools (see Fig 1B).

#### Exploitation of AMP computational prediction tools identifies >16,000 putative helminth AMP-LPs

Of the 64,581 helminth predicted peptides, peptides that were indicated as potential AMPs in >50% of the AMP computational prediction tools employed here (see Table 2) were assigned as antimicrobial peptide-like peptides (AMP-LPs), resulting in designation of 15,339 nematode and 1,501 flatworm-derived AMP-LPs (see Fig 1B and Fig 1Bi, ii). In order to prioritise the most promising helminth-derived AMP candidates from the total cohort, additional curation steps were applied to generate a dataset of higher-confidence helminth AMP-LPs.

#### Homology-based curation reduces the prioritised helminth AMP-LP cohort to 5,927 AMP-LPs

Of the 16,840 helminth AMP-LPs generated from the prediction phase of the pipeline, ~16% are representative of known helminth AMP groups [Cecropins, CSαβs, Diapausins, Nemapores and Glycine Rich Secreted Peptides; 19] and were removed from the pipeline (2,681 nematode AMP-LPs; see Table 5). Interestingly, for these previously identified putative nematode AMPs [19] the overall percentage of AMP prediction tools that predicted AMP activity varied (see Supplementary File 4). Indeed, for Cecropin peptides, with experimentally validated antimicrobial activity (39), the overall percentage of AMP prediction tools that predicted AMP activity ranged from 64-100%. These data highlight that, whilst the AMP prediction tools are a useful indicator of AMP potential, they should not independently direct putative AMP prioritisation.

**Table 5:** Traditional nematode AMP groups identified through the AMP prediction pipeline.

| Nematode AMP groups | No. of peptides identified | Overall prediction scores range |
| --- | --- | --- |
| Cecropin | 29 | 64-100% |
| Diapausin | 17 | 64-91% |
| CS$\alpha\beta$ peptides | 293 | 55-100% |
| Nemapore | 480 | 55-91% |
| Glycine Rich Secreted Peptides | 1862 | 55-100% |

Invertebrate AMPs often possess multiple homologs [50], such that the presence of AMP homologs within the helminth dataset would add further confidence in putative AMP designation. Therefore, the next step in the curation of the putative helminth AMP-LPs was the identification of AMP-LP sequelogs. Of the remaining 14,160 helminth AMP-LPs, 7,117 (50%) were assigned to 1,775 distinct groups based on their homology to each other. From this point, AMP-LPs were numbered according to their assigned group and designated nAMP-LP and fAMP-LP to distinguish between nematode and flatworm AMP-LPs respectively (see Supplementary Files 4 and 5). Putative helminth AMP-LPs that did not possess sequelogs were removed from the pipeline at this stage. Of the AMP-LP groups assigned based on homology, 84% encompassed <5 peptide members; this indicates limited putative AMP homology across helminth species (see Fig 1Biii). In addition, any AMP-LP groups that were only represented in a single genome were removed from the pipeline. Notably, as a result of this data filtering approach, species-specific AMP-LPs were unlikely to be prioritised, however all of the pre-filtered datasets remain available for future analysis.

#### Gene ontology-based data curation reveals 5,295 helminth AMP-LPs

Gene Ontology (GO) annotations were retrieved for the remaining helminth AMP-LPs. Any helminth AMP-LPs that were associated with non-AMP related Biological Process and Molecular Function GO terms were removed from the pipeline. It is possible that peptides with non-AMP GO annotations also possess antimicrobial activities, however this step was included to reduce the likelihood of false positives. This removed 525 nematode and 107 flatworm peptides from the prioritised helminth AMP-LP list, resulting in 5,295 helminth AMP-LPs.

For the helminth AMP-LPs that were removed, the associated non-AMP GO terms were analogous demonstrating that the AMP algorithms were comparably scoring sequences that possess similar functions (see Table 6). Many of these non-AMP GO annotations are associated with signalling processes or protease inhibition, such that it is unclear why these terms are commonly identified as antimicrobial by the AMP computational tools. It is possible that these peptides share characteristics of AMPs; indeed, many are cysteine-rich, a key feature of some AMP families [51]. Interestingly, in the flatworm dataset, many of the sequences with the ‘negative regulation of endopeptidase activity’ GO term (GO:0010951) possessed Kunitz-type inhibitor domains (Pfam: PF00014). Recently novel AMPs were identified within the C-terminus of the *S. mansoni* Kunitz Inhibitor protein (SmKI-1) [21]. This provides a rationale for the emergence of non-AMP GO associated peptides in the pipeline employed here and highlights the potential that some of the peptides removed at this stage of the pipeline may indeed contain regions which possess antimicrobial activity that could be explored in future.

**Table 6:** Common non-AMP related Gene Ontogeny (GO) terms associated with helminth AMP-LPs.

| Nematode |  |  |
| --- | --- | --- |
| *Five most common Biological Process GO terms* |  |  |
| GO accession | Name | % |
| GO:0007165 | Signal transduction | 18.9 |
| GO:0010951 | Negative regulation of endopeptidase activity | 12.6 |
| GO:0007218 | Neuropeptide signalling pathway | 11.6 |
| GO:0006644 | Phospholipid metabolic process | 9.8 |
| GO:0050482 | Arachidonic acid secretion | 9.8 |
| *Five most common Molecular Function GO terms* |  |  |
| GO accession | Name | % |
| GO:0004867 | Serine-type endopeptidase inhibitor activity | 20.3 |
| GO:0005179 | Hormone activity | 16.7 |
| GO:0004623 | Phospholipase A2 activity | 7.9 |
| GO:0003796 | Lysozyme activity | 4.8 |
| GO:0004531 | Deoxyribonuclease II activity | 2.9 |
| Flatworm |  |  |
| *Three most common Biological Process GO terms* |  |  |
| GO accession | Name | % |
| GO:0010951 | Negative regulation of endopeptidase activity | 60.3 |
| GO:0007165 | Signal transduction | 16.7 |
| GO:0006508 | Proteolysis | 10.3 |
| *Three most common Molecular Function GO terms* |  |  |
| GO accession | Name | % |
| GO:0004867 | Serine-type endopeptidase inhibitor activity | 70.6 |
| GO:0005179 | Hormone activity | 11.9 |
| GO:0008233 | Peptidase activity | 7.3 |

It is also interesting to note that some antimicrobial proteins such as lysozymes were also picked up in the pipeline described here. Although lysozymes were outside of the scope of this study as they are antimicrobial proteins [52], this pipeline identified shorter lysozyme sequences within the helminth predicted protein datasets. This indicated that the pipeline may also be useful in identifying shorter antimicrobial proteins or shorter antimicrobial fragments of longer proteins.

#### Curation of the helminth AMP-LPs based on peptide length resulted in a final dataset of 1,727 prioritised AMP-LPs

Most AMPs are <60 AAs in length; indeed, ~90% of invertebrate AMPs are <50 AAs [23]. As a result, a 60 AA length cut-off was implemented such that peptide groups, where the majority (>50%) of predicted peptides were >60AAs in length, were removed. This resulted in the removal of 3,568 peptides from a cohort of 5,295 leaving 1,727 putative helminth AMP-LPs in the final prioritised putative AMP cohort generated by the computational approach.

#### Cysteine-rich peptides dominate the prioritised helminth AMP-LPs

Cysteine-rich peptides are common across invertebrate AMP cohorts. Indeed, within helminths, three of the five known nematode AMP families are cysteine-rich [19]. Significantly, peptides with >2 cysteine residues (cysteine-rich) dominate the prioritised helminth AMP-LP cohort (72%). Cysteine-rich AMPs typically possess greater stability because of disulfide bond formation which aligns with their dominance in the helminth AMP datasets. Indeed, it was not surprising to find cysteine-rich peptides emerging from the pipeline as many of the computational platforms have been trained using cysteine-rich peptides. Due to the difficulties associated with production of peptides with >2 cysteine residues and the inability to verify correct disulfide bonding, cysteine-rich AMP-LPs were not explored further in this study.

#### Novel helminth AMP-LPs possess antibacterial activity

Of the 1,727 prioritised AMP-LPs identified from the computational AMP prediction pipeline, 479 AMP-LPs were predicted to have linear structure. Most of these (92%) originated from nematode species likely reflecting the expansion in genome datasets available for nematodes relative to flatworms. Twenty peptides (16 nematode, 4 flatworm) were selected for peptide synthesis based on: (i) peptide conservation across multiple species; (ii) overall AMP prediction tool scores; and (iii) evidence of expression in key helminth species and life stages. Due to helminth AMP-LP homology, the antimicrobial activity data arising from the 20 peptides screened here (see Table 7) informs the function of ~35% of the prioritised AMP-LP cohort.

**Table 7:**
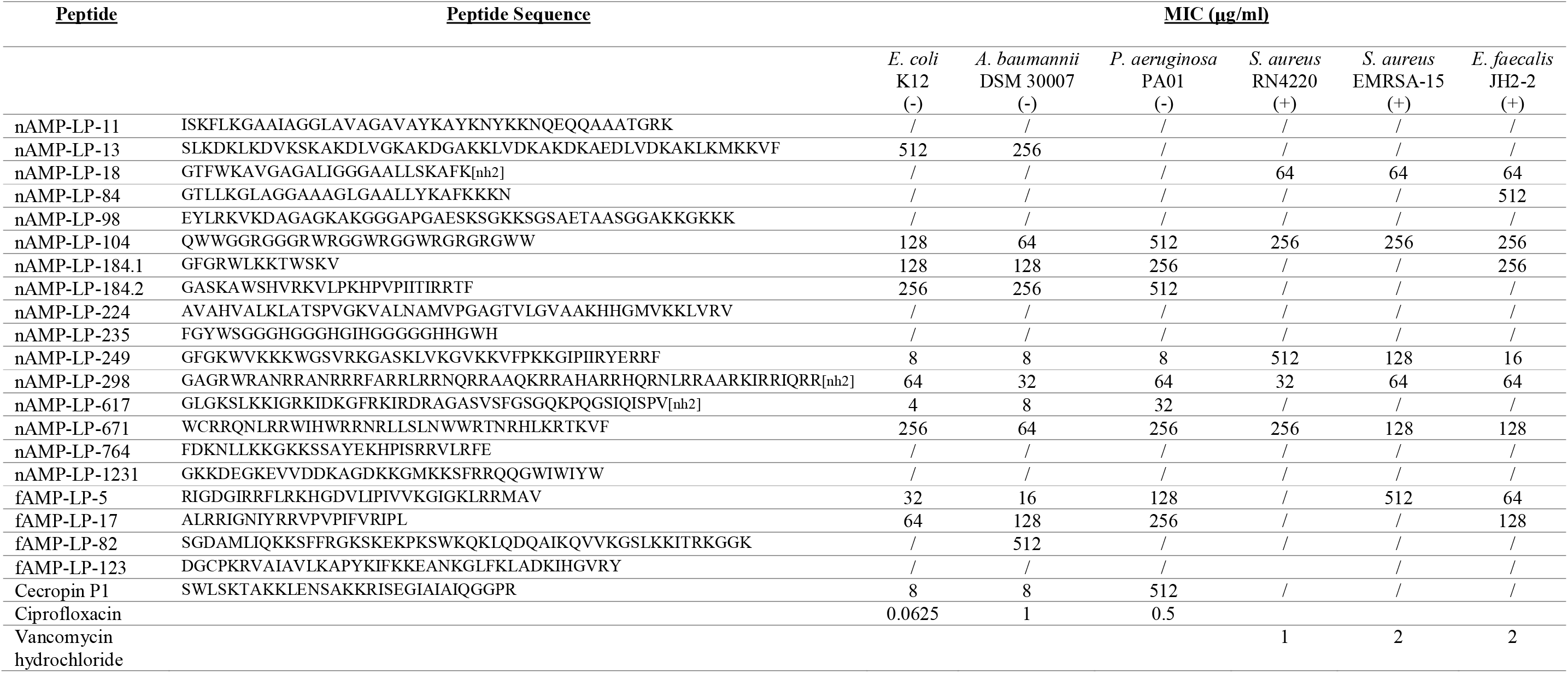
Minimum Inhibitory Concentrations (MICs) of selected helminth AMP-LPs against bacterial species.

All peptides were tested at >70% purity with the exception of Cecropin P1 which was used as a nematode control AMP at >95% purity. Ciprofloxacin was used as a positive control antibiotic for the gram-negative species (*Escherichia coli, Acinetobacter baumannii* and *Pseudomonas aeruginosa* and Vancomycin hydrochloride was used as a positive control antibiotic for the gram-positive species (*Staphylococcus aureus* and *Enterococcus faecalis*). / indicates that there was no activity at the highest concentration of peptide used (512μg/ml). Bacterial gram status is indicated by + or −. ‘n’ and ‘f’ before AMP-LP indicates a nematode and flatworm AMP-LP respectively.

Eight AMP-LPs displayed antibacterial activity (MIC < 100μg/ml) against at least one bacterial species (see Table 7). These data provide confidence that the integrated computational AMP prediction pipeline developed here can successfully identify novel bioactive AMPs. Significantly, most of the bioactive peptides displayed antibacterial activity against multiple species indicating broad-spectrum properties. AMPs can often have activities that are highly specific such that the absence of antimicrobial activity in 12 of the 20 AMP-LPs screened here does not rule out their potential as AMPs. Indeed, this may reflect antibacterial specificity for diverse pathogens that are not included in the screen employed in this study. Significantly, five helminth AMP-LPs (nAMP-LP-18, nAMP-LP-249, nAMP-LP-298, nAMP-LP-617, fAMP-LP-5) displayed notably potent and broad-spectrum antibacterial activities (see Table 7).

nAMP-LP-18 was identified in four *Meloidogyne* species (see Supplementary File 4). nAMP-LP-18 prepropeptides do not possess any known protein family domains. nAMP-LP-18 is highly conserved and located at the C-terminus of the precursor protein downstream to an additional peptide that is not predicted to be antimicrobial (see Fig 2A); note that multiple nAMP-LP-18-encoding genes are present in each *Meloidogyne* species (see Supplementary File 4). The synthesised nAMP-LP-18 (GTFWKAVGAGALIGGGAALLSKAFK-amide) displayed antibacterial activity against all of the gram-positive species tested but did not show activity against any of the gram-negative species screened (see Table 7). Based on a published life stage-specific *Meloidogyne incognita* RNAseq dataset [Study SRP109232, WBPSv16; 53] nAMP-LP-18-encoding genes (Minc3s01077g20499, Minc3s01204g21689, Minc3s07353g41022 and Minc3s09216g43021) are expressed in J3, J4 and adult female life stages (TPM>2). These data support the need to characterise the biological role and importance of these peptides in plant parasitic nematode species.

**Fig 2:**
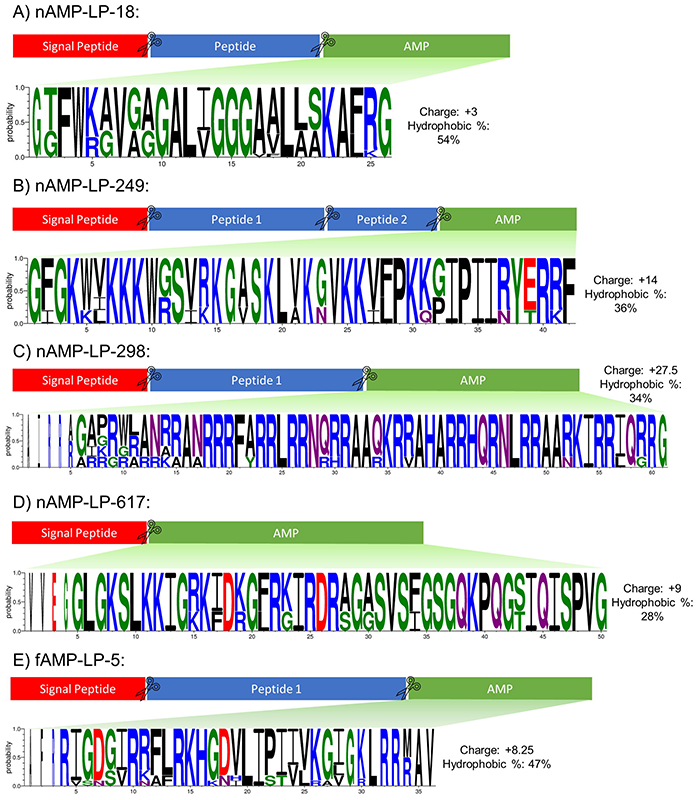
Protein precursor features and amino acid conservation of novel linear helminth AMPs. Signal peptide region indicated in red, predicted AMP region indicated in green and other peptides shown in blue. Scissors icon denotes region of putative cleavage sites. Note that boxes are not drawn to scale. Predicted AMP peptide conservation graphic shown below was generated using WebLogo3 [www.weblogo.threeplusone.com; 60, 61] using a multiple sequence alignment generated using Clustal Omega [29] of A) 21, B) 6, C) 5, D) 3, E) 11 peptides. WebLogo is coloured by amino acid chemistry [Green= Polar, Neutral= Purple, Basic= Blue, Acidic= Red, Hydrophobic= Black]. The height of amino acid letters indicates probability of an amino acid at a specific location. The consensus peptide sequence with gaps removed was used to obtain peptide charge and hydrophobic % using the APD3 Peptide Calculator and Predictor [https://aps.unmc.edu/prediction/predict; 23].

Multiple nAMP-LP-249-encoding genes were identified in *Trichuris suis* and *Trichuris trichuria* (see Supplementary File 4). nAMP-LP-249 prepropeptides do not possess any known protein family domains but two peptides, which were not predicted to have antimicrobial activity, are also encoded within the prepropeptide (see Fig 2B). The synthesised nAMP-LP-249 (GFGKWVKKKWGSVRKGASKLVKGVKKVFPKKGIPIIRYERRF) displayed potent activity against all gram-negative species screened (MIC 8μg/ml) as well as gram-positive *E. faecalis* (MIC 16μg/ml) but did not display activity against either *S. aureus* isolate (see Table 7). The potent broad-spectrum activity displayed by this peptide highlights the need to expand activity screens towards a more diverse portfolio of microbes.

nAMP-LP-298 is highly conserved across four Clade 9/V gastrointestinal parasites (*Ancylostoma ceylanicum*, *Haemonchus contortus*, *Haemonchus placei* and *Teladorsagia circumcincta* (see Supplementary File 4). nAMP-LP-298 is rich in arginine residues resulting in a net charge of +27.5 (see Fig 2C). While most known AMPs have a positive net charge [88% of AMPs on the APD3 have a net positive charge; 23] only 4 AMPs in the APD3 have a net charge >20 (AP00411, AP00684, AP02232 and AP03367). Cationic AMPs are generally considered to interact with negatively charged bacterial membranes through electrostatic interactions [54] and, for some modified AMPs, increasing the peptide charge has resulted in more potent antimicrobial activity [55-57]. nAMP-LP-298 (GAGRWRANRRANRRRFARRLRRNQRRAAQKRRAHARRHQRNLRRAARKIRRIQRR-amide; see Table 7) was only peptide tested in this study that displays activity against all bacterial species screened; this may be due to the high peptide charge which could be unravelled through further work.

nAMP-LP-617 was only identified in the rodent parasite *Heligmosomoides polygyrus* indicating that these peptides are highly restricted in nematodes (see Supplementary File 4). The nAMP-LP-617 prepropeptides does not encode any known protein family domains and nAMP-LP-617 appears to be the only peptide encoded on the genes (see Fig 2D). The tested nAMP-LP-617 (GLGKSLKKIGRKIDKGFRKIRDRAGASVSFGSGQKPQGSIQISPV-amide) displayed potent activity against gram-negative species (MIC 4-32μg/ml; see Table 7) but was not active against gram-positive species. Based on a published *H. polygyrus* RNAseq study [SRP157940, WBPSv16; 58] all three genes (HPOL_0001299301, HPOL_0001299401 and HPOL_0002106401) appear to be expressed (TPM >2).

fAMP-LP-5 was identified in three *Taenia* species (see Supplementary File 5). fAMP-LP-5 prepropeptides do not display any known protein family domains but appear to encode another peptide which was not predicted to have antimicrobial activity (see Fig 2E). The tested fAMP-LP-5 (RIGDGIRRFLRKHGDVLIPIVVKGIGKLRRMAV) displayed some selective gram-negative activity (againt *E. coli* and *A. baumannii*) and gram-positive activity (*E. faecalis*) (see Table 7). This peptide represents the first AMP to be identified from cestode species.

#### AMP-LPs are present in helminth biofluids validating computational AMP-LP identification

To validate the computational AMP prediction pipeline developed here the presence of AMP-LPs in nematode biofluids was examined. LC-MS/MS analysis of *A. suum* pseudocoelomic fluid (*As-*PCF) detected 60 high-confidence peptide spectrum matches (PSMs) above the 1% false discovery rate (FDR) threshold (−10lgP 47.3, Supplementary File 6A-C). The 60 PSMs correlate to 15 AMP-LPs detected via the computational AMP prediction pipeline described here, nine of which were detected within the predicted mature peptide region (see Table 8, Fig 3 and Supplementary File 6). Of these nine AMP-LPs, three were identified in 100% of *As*-PCF biological replicates, one was identified in 67% of *As*-PCF biological replicates and the remaining five were detected in at least one biological replicate (see Table 8 and Supplementary File 6 and 7). AMP-LP-835 was detected with the highest confidence and AMP-LP-2454 was detected with the highest level of coverage (see Table 8, Supplementary File 6). Reduction of PSM cut-offs (P-value <0.05) enhanced peptide sequence coverage (see Table 8 and Fig 3); for example, AMP-LP-2454 sequence coverage increases to from 30% to 92% when PSM cut-off is reduced from 1% FDR to P-value <0.05 (see Table 8 and Fig 3). In addition to enhanced sequence coverage, the number of PSMs also increased from 60 (1% FDR) to 85 and 127 when the cut-off was reduced to P-value <0.01 and <0.05, respectively. This highlights a need for further discussion of the application of FDR cut-off ranges in peptidomics analysis [59]. The LC-MS/MS analysis performed here provides the first experimental evidence for the presence of novel nematode-derived AMP-LPs in helminth biofluid and validates the application of the computational AMP prediction pipeline for novel AMP discovery which could be readily translated to other invertebrate species.

**Table 8.**
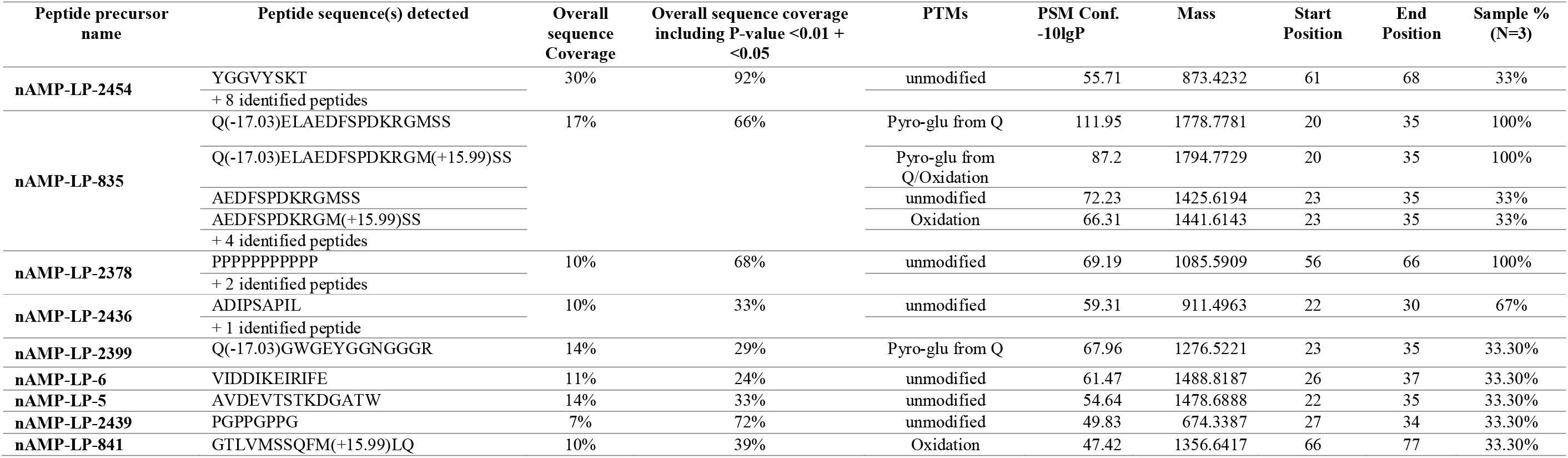
Summary of detected nAMP-LPs identified in pooled female *Ascaris suum* pseudocoelomic fluid via LC-MS/MS.

**Fig 3:**
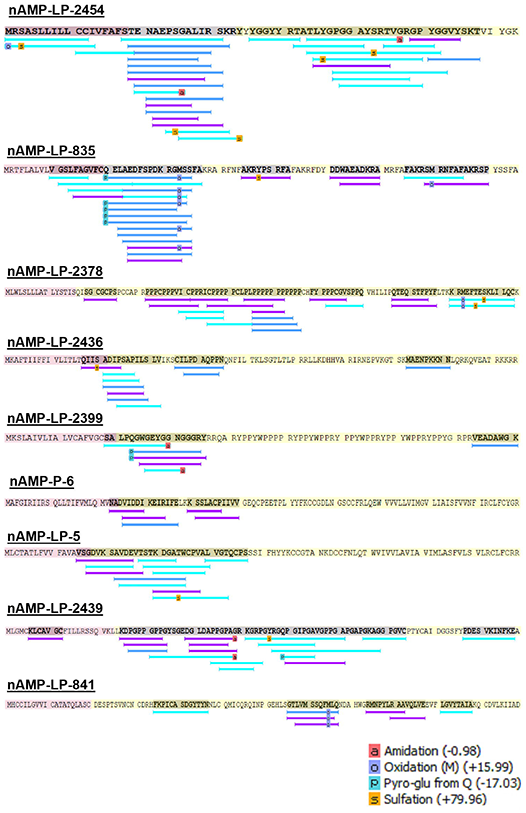
Peptide spectrum matches (PSMs) for each nAMP-LP detected via LC-MS/MS methods. Peptides were only included here if PSMs were detected at FDR 1% within the mature peptide region. Red shading indicates the signal peptide, yellow shading indicates the predicted mature peptide region. Bars underneath the sequence indicate the presence of a PSM. Dark blue shaded bars indicate a PSM found at 1% FDR (−10lgP score > 47.3), purple shaded bars indicate a PSM found > P-value 0.05 cut off (Peptide-10lgP score >20) but below the 1% FDR cut-off, light blue shading indicates a PSM found > P-value 0.01 cut-off (Peptide-10lgP score >13) but below the P-value 0.05 cut off-10lgP score of 20. All relevant spectra for each can be found in Supplementary File 7.

Only sequences of the putative mature nAMP-LPs are shown. Peptide sequences detected relate to sequences detected to a 1% FDR via PEAKS studio X (Bioinformatics solution Inc., Waterloo, ON, Canada). Additional identified peptide data consists of overlapping peptide spectrum match sequences detected for each peptide including propeptides belonging to the nAMP-LP precursors and relevant data for each can be viewed in Supplementary File 6. Overall sequence coverage denotes the % of the prepropeptide nAMP-LP sequence detected via LC-MS/MS and includes additional identified peptides. Overall sequence coverage including P-value <0.01 + <0.05 denotes the % of prepropeptide nAMP-LP sequences detected via LC-MS/MS when PSM cut-offs are lowered to P-value <0.01 and <0.05 (see Supplementary File 6). PTMs; Post-Translational Modifications. PSM Conf.-10lgP; Peptide Spectrum Match confidence score determined by PEAKS studio X and converted to-10lgP. Start Position and End position relate to the position in the prepropeptide nAMP-LP sequence to which the PSM relates to. Sample % denotes the percentage of samples peptides were detected in.

## Conclusions

This study uncovers AMP diversity in helminths through the integration of novel computational AMP discovery approaches on a pan phylum scale. The data demonstrate that flatworms do not encode traditional lophotrochozoan AMP groups and underscores the need for innovative approaches to AMP discovery across helminth phyla. Here we report the development and application of a novel integrated computational AMP discovery pipeline to reveal >16,000 peptides with antimicrobial potential from 127 helminth species. These data represent a unique database of helminth-derived putative AMPs ripe for therapeutic exploitation. Indeed, several of the novel AMP-LPs identified in this study have potent antibacterial activities against relevant bacterial pathogens of humans and animals. In addition, helminth AMP-LPs are present in nematode biofluids providing the drive to better understand the importance of AMPs to helminth biology and the host-worm-microbiome relationship that will support drug discovery programmes for helminth parasites.

## Acknowledgements

The authors thank Dr Linda Oyama for antimicrobial assay assistance and Karro, Cookstown, NI for help with the collection of *Ascaris suum*.

## Supporting information

**S1 File. Lophotrochozoan-derived AMPs identified from literature search**.

**S2 File. Genomic datasets utilised in the flatworm homology approaches**.

**S3 File. Genomic datasets utilised in the helminth AMP prediction approach. S4 File. Nematode AMP-LPs**.

**S5 File. Flatworm AMP-LPs**.

**S6 File. nAMP-LPs detected in pooled female As-PCF via LC-MS/MS**

**S7 File. All peptide spectrum match MS/MS spectra detected for each nAMP-LP found in Fig 3**.

## Notes

### Competing Interest Statement

The authors have declared no competing interest.

## References

1. Zasloff M. Antimicrobial Peptides of Multicellular Organisms: My Perspective. In: Matsuzaki K, editor. Antimicrobial Peptides. Springer Singapore; 2019. pp. 3–6.

2. Magana M, Pushpanathan M, Santos AL, Leanse L, Fernandez M, Ioannidis A, et al. The value of antimicrobial peptides in the age of resistance. Lancet Infect Dis. 2020;20(9): e216–e230.

3. Wang G. Prediction and Design of Antimicrobial Peptides: Methods and Applications to Genomes and Proteomes. In: Wang G, editor. Antimicrobial Peptides: Discovery, Design and Novel Therapeutic Strategies 2^nd^ edition. CAB International; 2017. pp. 101–118.

4. Wang G, Zietz CM, Mudgapalli A, Wang S, Wang Z. The evolution of the antimicrobial peptide database over 18 years: Milestones and new features. Protein Sci. 2021;31(1): 92–106.

5. Wu Q, Ke H, Li D, Wang Q, Fang J, Zhou J. Recent Progress in Machine Learning-based Prediction of Peptide Activity for Drug Discovery. Curr Top Med Chem. 2019;19(1): 4–16.

6. Melo MCR, Maasch JRMA, de la Fuente-Nunez C. Accelerating antibiotic discovery through artificial intelligence. Commun Biol. 2021;4(1): 1050.

7. Benoist L, Houyvet B, Henry J, Corre E, Zanuttini B, Zatylny-Gaudin C. In-Depth In Silico Search for Cuttlefish (*Sepia officinalis*) Antimicrobial Peptides Following Bacterial Challenge of Haemocytes. Mar Drugs. 2020;18(9): 439.

8. Moretta A, Salvia R, Scieuzo C, Di Somma A, Vogel H, Pucci P, et al. A bioinformatic study of antimicrobial peptides identified in the Black Soldier Fly (BSF) *Hermetia illucens* (Diptera: Stratiomyidae). Sci Rep. 2020;10(1): 16875.

9. Ma Y, Guo Z, Xia B, Zhang Y, Liu X, Yu Y, et al. Identification of antimicrobial peptides from the human gut microbiome using deep learning. Nat Biotechnol. 2022;40: 1–11.

10. Wegener Parfrey L, Jirků M, Šíma R, Jalovecka M, Sak B, Grigore K, et al. A benign helminth alters the host immune system and the gut microbiota in a rat model system. PLoS One. 2017;12(8): e0182205.

11. Slater R, Frau A, Hodgkinson J, Archer D, Probert C. A Comparison of the Colonic Microbiome and Volatile Organic Compound Metabolome of *Anoplocephala perfoliata* Infected and Non-Infected Horses: A Pilot Study. Animals. 2021;11(3): 755.

12. Xu M, Jiang Z, Huang W, Yin J, Ou S, Jiang Y, et al. Altered Gut Microbiota Composition in Subjects Infected With *Clonorchis sinensis*. Front Microbiol. 2018;9: 2292.

13. Portet A, Toulza E, Lokmer A, Huot C, Duval D, Galinier R, et al. Experimental Infection of the *Biomphalaria glabrata* Vector Snail by *Schistosoma mansoni* Parasites Drives Snail Microbiota Dysbiosis. Microorganisms. 2021;9(5): 1084.

14. Jenkins TP, Pritchard DI, Tanasescu R, Telford G, Papaiakovou M, Scotti R, et al. Experimental infection with the hookworm, *Necator americanus*, is associated with stable gut microbial diversity in human volunteers with relapsing multiple sclerosis. BMC Biol. 2021;19(1): 74.

15. Gobert GN, Atkinson LE, Lokko A, Yoonuan T, Phuphisut O, Poodeepiyasawat A, et al. Clinical helminth infections alter host gut and saliva microbiota. PLoS Negl Trop Dis. 2022;16(6): e0010491.

16. Marra A, Hanson MA, Kondo S, Erkosar B, Lemaitre B. *Drosophila* antimicrobial peptides and lysozymes regulate gut microbiota composition and abundance. Mbio. 2021;12(4): e00824–21.

17. Fraune S, Augustin R, Anton-Erxleben F, Wittlieb J, Gelhaus C, Klimovich VB, et al. In an early branching metazoan, bacterial colonization of the embryo is controlled by maternal antimicrobial peptides. Proc Natl Acad Sci U.S.A. 2010;107(42): 18067–18072.

18. Tarr DEK. Distribution and characteristics of ABFs, cecropins, nemapores, and lysozymes in nematodes. Dev Comp Immunol. 2012;36(3): 502–520.

19. Irvine A, Huws SA, Atkinson LE, Mousley A. The Nematode Antimicrobial Peptidome: a novel opportunity for parasite control? BioRxiv [Preprint] 2022 bioRxiv: 507570 [Preprint]. 2022 [cited 2022 December 16]. Available from: https://www.biorxiv.org/content/10.1101/2022.09.26.507570v1 doi: 10.1101/2022.09.26.507570.

20. Quinn GAP, Heymans R, Rondaj F, Shaw C, de Jong-Brink M. *Schistosoma mansoni* dermaseptin-like peptide: structural and functional characterization. J Parasitol. 2005;91(6): 1340–1351.

21. Santos BPO, Alves ESF, Ferreira CS, Ferreira-Silva A, Góes-Neto A, Verly RM, et al. Schistocins: Novel antimicrobial peptides encrypted in the *Schistosoma mansoni* Kunitz Inhibitor SmKI-1. Biochim Biophys Acta Gen Subj. 2021;1865(11): 129989.

22. Bleidorn C. Recent progress in reconstructing lophotrochozoan (spiralian) phylogeny. Org Divers Evol. 2019;19(4): 557–566.

23. Wang G, Li X, Wang Z. APD3: the antimicrobial peptide database as a tool for research and education. Nucleic Acids Res. 2015;44(D1): D1087–D1093.

24. Waghu FH, Idicula◻Thomas S. Collection of antimicrobial peptides database and its derivatives: Applications and beyond. Protein Sci. 2019;29(1): 36–42.

25. Piotto SP, Sessa L, Concilio S, Iannelli P. YADAMP: yet another database of antimicrobial peptides. Int J Antimicrob Agents. 2012;39(4): 346–351.

26. Pirtskhalava M, Amstrong AA, Grigolava M, Chubinidze M, Alimbarashvili E, Vishnepolsky B, et al. DBAASP v3: database of antimicrobial/cytotoxic activity and structure of peptides as a resource for development of new therapeutics. Nucleic Acids Res. 2021;49(D1): D288–D297.

27. Gomez EA, Giraldo P, Orduz S. InverPep: A database of invertebrate antimicrobial peptides. J Glob Antimicrob Resist. 2017;8: 13–17.

28. Shi G, Kang X, Dong F, Liu Y, Zhu N, Hu Y, et al. DRAMP 3.0: an enhanced comprehensive data repository of antimicrobial peptides. Nucleic Acids Res. 2022;50(D1): D488–D496.

29. Madeira F, Pearce M, Tivey ARN, Basutkar P, Lee J, Edbali O, et al. Search and sequence analysis tools services from EMBL-EBI in 2022. Nucleic Acids Res. 2022;50(W1): W276–W279.

30. Howe KL, Bolt BJ, Shafie M, Kersey P, Berriman M. WormBase ParaSite− a comprehensive resource for helminth genomics. Mol. Biochem. Parasitol. 2017;215: 2–10.

31. Nielsen H. Predicting secretory proteins with SignalP. In: Kihara D, editor. Protein function prediction: Methods and Protocols. Humana New York; 2017. pp. 59–73.

32. Duckert P, Brunak S, Blom N. Prediction of proprotein convertase cleavage sites. Protein Eng Des Sel. 2004;17(1): 107–112.

33. Lee HT, Lee CC, Yang JR, Lai JZC, Chang KY. A Large-Scale Structural Classification of Antimicrobial Peptides. BioMed Res Int. 2015;475062.

34. Xiao X, Wang P, Lin WZ, Jia JH, Chou KC. iAMP-2L: A two-level multi-label classifier for identifying antimicrobial peptides and their functional types. Anal Biochem. 2013;436(2): 168–177.

35. Meher PK, Sahu TK, Saini V, Rao AR. Predicting antimicrobial peptides with improved accuracy by incorporating the compositional, physico-chemical and structural features into Chou’s general PseAAC. Sci Rep. 2017;7: 42362.

36. Veltri D, Kamath U, Shehu A. Deep learning improves antimicrobial peptide recognition. Bioinformatics. 2018;34(16): 2740–2747.

37. Bhadra P, Yan J, Li J, Fong S, Siu SWI. AmPEP: Sequence-based prediction of antimicrobial peptides using distribution patterns of amino acid properties and random forest. Sci Rep. 2018;8: 1697.

38. Ashburner M, Ball CA, Blake JA, Botstein D, Butler H, Cherry JM, et al. Gene Ontology: tool for the unification of biology. Nat Genet. 2000;25: 25–29.

39. The Gene Ontology Consortium. The Gene Ontology resource: enriching a GOld mine. Nucleic Acids Res. 2021;49(D1): D325–D334.

40. Blum M, Chang H-Y, Chuguransky S, Grego T, Kandasaamy S, Mitchell A, et al. The InterPro protein families and domains database: 20 years on. Nucleic Acids Res. 2021;49(D1): D344–D354.

41. Mistry J, Chuguransky S, Williams L, Qureshi M, Salazar GA, Sonnhammer ELL, et al. Pfam: The protein families database in 2021. Nucleic Acids Res. 2021;49(D1): D412–D419.

42. Wagner GP, Kin K, Lynch VJ. A model based criterion for gene expression calls using RNA-seq data. Theory Biosci. 2013;132: 159–164.

43. Oyama LB, Girdwood SE, Cookson AR, Fernandez-Fuentes N, Privé F, Vallin HE, et al. The rumen microbiome: an underexplored resource for novel antimicrobial discovery. NPJ Biofilms Microbiomes. 2017;3: 33.

44. Onime LA, Oyama LB, Thomas BJ, Gani J, Alexander P, Waddams KE, et al. The rumen eukaryotome is a source of novel antimicrobial peptides with therapeutic potential. BMC Microbiol. 2021;21: 105.

45. Atkinson LE, Liu Y, McKay F, Vandewyer E, Viau C, Irvine A, et al. *Ascaris suum* Informs Extrasynaptic Volume Transmission in Nematodes. ACS Chem Neurosci. 2021;12: 3176–3188.

46. Rappsilber J, Mann M, Ishihama Y. Protocol for micro-purification, enrichment, pre-fractionation and storage of peptides for proteomics using StageTips. Nat. Protoc. 2007;2: 1896–1906.

47. Tarr DEK. Establishing a reference array for the CS-αβ superfamily of defensive peptides. BMC Res Notes. 2016;9: 490.

48. Gerdol M, Schmitt P, Venier P, Rocha G, Rosa RD, Destoumieux-Garzón D. Functional Insights From the Evolutionary Diversification of Big Defensins. Front Immunol. 2020;11: 758.

49. The UniProt Consortium. UniProt: the universal protein knowledgebase in 2021. Nucleic Acids Res. 2021;49(D1): D480–D489.

50. Tassanakajon A, Somboonwiwat K, Amparyup P. Sequence diversity and evolution of antimicrobial peptides in invertebrates. Dev Comp Immunol. 2015;48(2): 324–341.

51. Lehrer RI. Evolution of Antimicrobial Peptides: A View from the Cystine Chapel. In: Hiemstra PS, Zaat SAJ, editors. Antimicrobial Peptides and Innate Immunity: Springer Basel; 2013. pp. 1–27.

52. Ragland SA, Criss AK. From bacterial killing to immune modulation: Recent insights into the functions of lysozyme. PLOS Pathog. 2017;13(9): e1006512.

53. Choi I, Subramanian P, Shim D, Oh B-J, Hahn B-S. RNA-Seq of plant-parasitic nematode *Meloidogyne incognita* at various stages of its development. Front. Genet. 2017;8: 190.

54. Ciumac D, Gong H, Hu X, Lu JR. Membrane targeting cationic antimicrobial peptides. J. Colloid Interface Sci. 2019;537: 163–185.

55. Xie J, Zhao Q, Li S, Yan Z, Li J, Li Y, et al. Novel antimicrobial peptide CPF[C1 analogs with superior stabilities and activities against multidrug[resistant bacteria. Chem Biol Drug Des. 2017;90(5): 690–702.

56. Zhou J, Liu Y, Shen T, Chen L, Zhang C, Cai K, et al. Antimicrobial activity of the antibacterial peptide PMAP-36 and its analogues. Microb Pathog. 2019;136: 103712.

57. Mangmee S, Reamtong O, Kalambaheti T, Roytrakul S, Sonthayanon P. Antimicrobial Peptide Modifications against Clinically Isolated Antibiotic-Resistant *Salmonella*. Molecules. 2021;26(15): 4654.

58. Rausch S, Midha A, Kuhring M, Affinass N, Radonic A, Kühl AA, et al. Parasitic Nematodes Exert Antimicrobial Activity and Benefit From Microbiota-Driven Support for Host Immune Regulation. Front. Immunol. 2018;9: 2282.

59. Foreman RE, George AL, Reimann F, Gribble FM, Kay RG. Peptidomics: A Review of Clinical Applications and Methodologies. 2021;20(8): 3782–3797.

60. Schneider TD, Stephens RM. Sequence logos: a new way to display consensus sequences. Nucleic Acids Res. 1990;18(20): 6097–6100.

61. Crooks GE, Hon G, Chandonia J-M, Brenner SE. WebLogo: A Sequence Logo Generator. Genome Res. 2004;14: 1188–1190.

